## Supplemental Information for "Regularized Single-cell Imaging Enables Generalizable AI models for Stain-free Cell Viability Screening"

**Short title:** Regularized Imaging for Generalizable AI

**Authors:** Pan Deng[1,2], Deasung Jang[1,2], Samuel G. Berryman[1,2], Simon P. Duffy[2,3], and Hongshen Ma[1,2,4,5]* (ORCID: 0000-0001-5459-6493)

**Affiliations**
[1]Department of Mechanical Engineering, University of British Columbia
[2]Centre for Blood Research, University of British Columbia
[3]British Columbia Institute of Technology
[4]School of Biomedical Engineering, University of British Columbia
[5]Vancouver Prostate Centre, Vancouver General Hospital

**Corresponding Author**
Hongshen Ma
2054-6250 Applied Science Lane
Vancouver, BC, Canada V6T 1Z4
604.722.5382


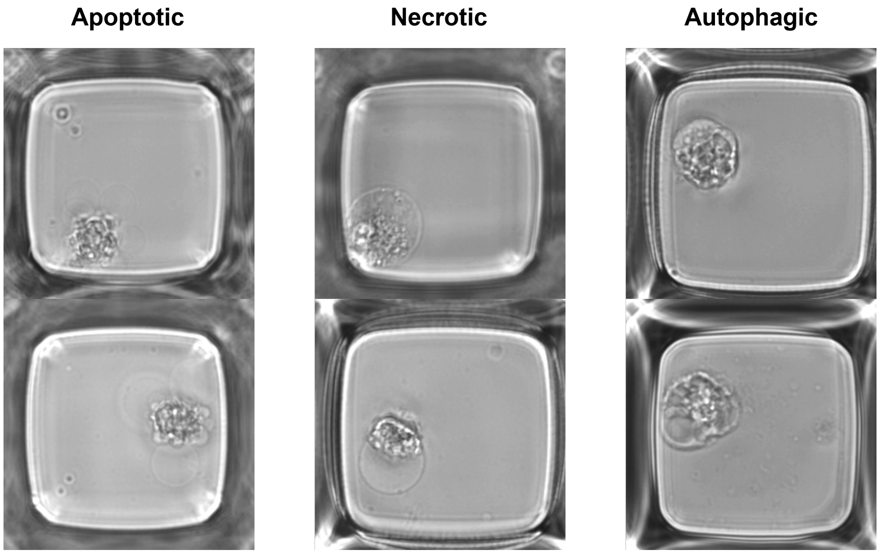


**Fig. S1**. Representative images of apoptotic cells, necrotic cells and autophagic cells in the training dataset.

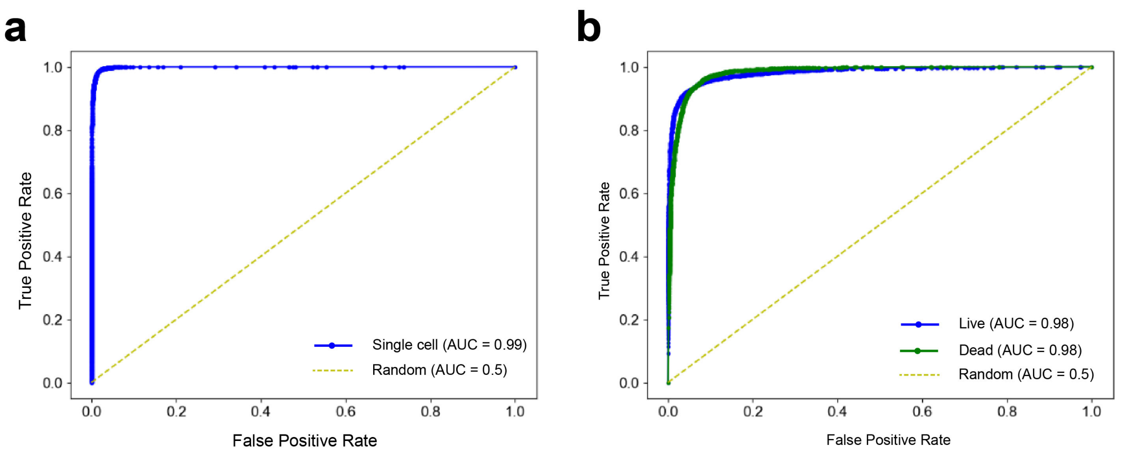


**Fig. S2**. The ROC graphs for the CNN model to identify nanowells occupied by a single cell (**a**) and the CNN model to assess the live/dead status of single cells (**b**). The diagonal dashed line indicates a random guess with an AUC of 0.5.

| | **Condition** | **Death mechanisms** | Ref. |
|---|---|---|---|
| **Training dataset** | Complete culture media | Apoptosis | [1] |
| | Culture media – no FBS | Apoptosis<br>Autophagy | [2] |
| | Ethanol | Apoptosis<br>Necrosis<br>Autophagy | [3] |
| | Andrographolide | Apoptosis<br>Autophagy | [4,5] |
| | Daunorubicin | Apoptosis | [6,7] |
| **Testing dataset** | Bortezomib | Apoptosis | [8] |
| | Cisplatin | Apoptosis<br>Necrosis | [9,10] |
| | Staurosporine | Apoptosis | [11,12] |
| | Doxorubicin | Apoptosis | [13,14] |
| | Carfilzomib | Apoptosis<br>Necrosis<br>Autophagy | [14–16] |
| | Mitoxantrone | Apoptosis<br>Autophagy | [17,18] |
| | Epirubicin | Apoptosis | [19] |
| | Aprepitant | Apoptosis | [20] |

**Table S1.** Cell death mechanisms in different conditions.
